# Structural insights into VRC01-class bnAb precursors with diverse light chains elicited in the IAVI G001 human vaccine trial

**DOI:** 10.1101/2025.05.22.655646

**Authors:** Xiaohe Lin, Christopher A. Cottrell, Oleksandr Kalyuzhniy, Ryan Tingle, Michael Kubitz, Danny Lu, Meng Yuan, William R. Schief, Ian A. Wilson

**Affiliations:** Department of Integrative Structural and Computational Biology, The Scripps Research Institute, La Jolla, California, USA; Center for HIV/AIDS Vaccine Development, The Scripps Research Institute, La Jolla, CA 92037, USA; Department of Immunology and Microbial Science, The Scripps Research Institute, La Jolla, CA 92037, USA; IAVI Neutralizing Antibody Center, The Scripps Research Institute, La Jolla, CA 92037, USA; Moderna Inc., Cambridge, MA, 02139, USA

## Abstract

The development of germline-targeting vaccines represents a potentially transformative strategy to elicit broadly neutralizing antibodies (bnAbs) against HIV and other antigenically diverse pathogens. Here, we report on structural characterization of vaccine-elicited VRC01-class bnAb precursors in the IAVI G001 Phase 1 clinical trial with the eOD-GT8 60mer nanoparticle as immunogen. High-resolution X-ray structures of eOD-GT8 monomer complexed with Fabs of five VRC01-class bnAb precursors with >90% germline identity revealed a conserved mode of binding to the HIV CD4-binding site (CD4bs) via IGHV1-2-encoded heavy chains, mirroring mature bnAb interactions. The light-chain V-gene diversity emulated VRC01 bnAbs and stabilized antigen engagement, while their conserved five-residue LCDR3 motifs prevented steric clashes. Notably, the VRC01-class bnAb precursors accommodated the N276 glycan, a key barrier in HIV Env recognition, through structural rearrangements in HCDR3 or LCDR1, despite its absence in the immunogen. Surface Plasmon Resonance (SPR) analysis showed that 87% of elicited antibodies retained glycan binding capacity, albeit with reduced affinity. These findings validate the ability of eOD-GT8 60mer nanoparticles to prime VRC01-class bnAb precursors with native-like paratopes but with intrinsic glycan adaptability. Structural mimicry of mature bnAbs was observed even with limited somatic hypermutation, indicating that critical features are encoded in the germline repertoire. The structures highlight how germline-encoded features drive bnAb-like recognition at early stages. This work provides molecular evidence supporting germline-targeting in humans and provides guidance for designing booster immunogens to shepherd affinity maturation toward broad neutralization.

## Introduction

Vaccine elicitation of bnAbs, known for their ability to target conserved epitopes on the HIV envelope glycoprotein (Env), represents a promising avenue for achieving long-lasting protection against diverse HIV-1 viral strains (1, 2). However, the rarity of human naive B cells with bnAb-precursor properties and the low affinities of such naive precursors for native Env proteins have hindered traditional vaccine approaches (3-5). The germline-targeting vaccine strategy has emerged as a potentially transformative approach in the field of antibody research, aiming to overcome critical challenges in eliciting bnAbs against human pathogens with high antigenic diversity, such as HIV (2, 6-8).

Germline-targeting vaccine design addresses this challenge in part by engineering priming immunogens that bind with high specificity and affinity to germline gene-encoded precursors of bnAbs (4, 5, 9-24). These germline-targeting priming immunogens aim to recruit and activate rare bnAb-precursor B cells, priming them for subsequent maturation into potent neutralizers through booster immunizations (13, 14, 21, 25-31). The engineered outer domain (eOD) germline-targeting version 8 (eOD-GT8), which was designed to activate VRC01-class germline-precursor B cells specific for the CD4bs on HIV Env, exemplifies this strategy (4, 9-12, 19, 25, 26, 30, 31). VRC01-class B cell receptors are defined by their specificity for the CD4bs and by using heavy chains derived from variable (V) gene alleles IGHV1-2*02 or *04 paired with light chains containing a five-amino-acid LCDR3 (4, 5, 32, 33). The eOD-GT8 has been displayed on a multivalent 60-mer nanoparticle to enhance B cell receptor engagement and facilitate germinal center (GC) responses, and the resulting nanoparticle has been termed eOD-GT8 60mer (4, 9-12, 19, 30, 31).

The IAVI G001 phase 1 clinical trial marked a pivotal milestone in testing germline-targeting vaccine strategies in humans. Conducted on healthy adult volunteers, the trial evaluated the safety, tolerability, and immunogenicity of the eOD-GT8 60mer immunogen adjuvanted with AS01_B_ (19, 34, 35). A key objective of the trial was to determine whether eOD-GT8 60mer could effectively prime VRC01-class B cell responses in humans, as it had in preclinical models (4, 9-11, 19, 30, 31). Results from the trial demonstrated that the immunogen successfully activated VRC01-class bnAb precursors in 97% of vaccine recipients and induced somatic hypermutation (SHM), signaling a critical first step toward the end goal of maturation into bnAbs. Notwithstanding, the molecular mechanisms that enabled germline precursors to engage the immunogen and their structural compatibility with mature bnAb binding modes has remained uncharacterized. Additionally, the capacity of these early elicited antibodies (Abs) to adapt to conserved glycans in the extensive glycan shield surrounding the Env protein (36, 37), such as the N276 glycan (27, 38), has not yet been resolved.

This study addresses these gaps through high-resolution structural characterization of IAVI G001 (hereafter referred to as G001) vaccine-elicited antibodies in complex with eOD-GT8. By resolving crystal structures of antibody Fab fragments bound to the immunogen, we were able to define how germline-encoded features in the elicited bnAb-precursor antibodies mediate epitope recognition and identify structural parallels between VRC01-class bnAb precursors and mature bnAbs. We further investigate the role of light-chain diversity in stabilizing antigen interactions and evaluate glycan accommodation mechanisms critical for guiding affinity maturation. These findings further validate the germline-targeting strategy and provide guidance for designing booster immunogens to steer antibody precursor evolution toward broad neutralization (25-28, 30, 31).

## Results

### X-ray crystal structures of eOD-GT8 complexes

To investigate how VRC01-class bnAb precursors elicited in G001 interact structurally with the eOD-GT8 antigen, we selected five VRC01-class Abs from the trial (G001-0087, G001-58, G001-59, G001-179, and G001-14) for crystallographic analysis of eOD-GT8 complexes. The selected Abs were broadly representative of the VRC01-class Abs isolated in IAVI G001, in that they: (i) were derived from different types of samples (plasmablasts, memory B cells, and GC B cells) collected from multiple participants at different time points; (ii) utilized V_H_1-2*02 or V_H_1-2*04 heavy-chain alleles, both of which were observed among G001 VRC01-class bnAb precursors (19, 35); (iii) utilized the four most common light chain variable genes observed among VRC01-class bnAb precursors in G001: IGHV1-33, IGHV3-20, IGHV1-5, and IGHV3-15, genes that are also utilized by VRC01-class bnAbs (19); (iv) had modest degrees of SHM consistent with that reported for G001 (i.e. the selected Abs were not SHM outliers); (v) had no insertions or deletions, consistent with the vast majority of VRC01-class responses in G001; and (vi) had affinities, on-rates, and off-rates for eOD-GT8 consistent with the ranges reported for VRC01-class responses in G001 (Table S1) (19). Two versions of eOD-GT8 were used: eOD-GT8-mingly, which represents a minimal glycan variant of the antigen used in the trial, and eOD-GT8-mingly-N276, which includes the conserved N276 glycan so as to evaluate its accommodation by the VRC01-class bnAb precursors.

We determined crystal structures for all five G001 trial-derived VRC01-class antibody Fab fragments (G001-0087, G001-58, G001-59, G001-179, and G001-14) in complex with eOD-GT8-mingly or eOD-GT8-mingly-N276 (Fig. 1A, Table S2). Structural analysis revealed that all five antibodies engage the CD4bs with similar binding mode to that of VRC01-class bnAbs (33, 39-46), (Fig. 1B). The Cα RMSD between the Fab variable domain of each trial-derived antibody and bnAb VRC01 is 0.73 Å, 0.89 Å, 0.82 Å, 0.80 Å, and 0.78 Å, respectively.

**Fig. 1.**
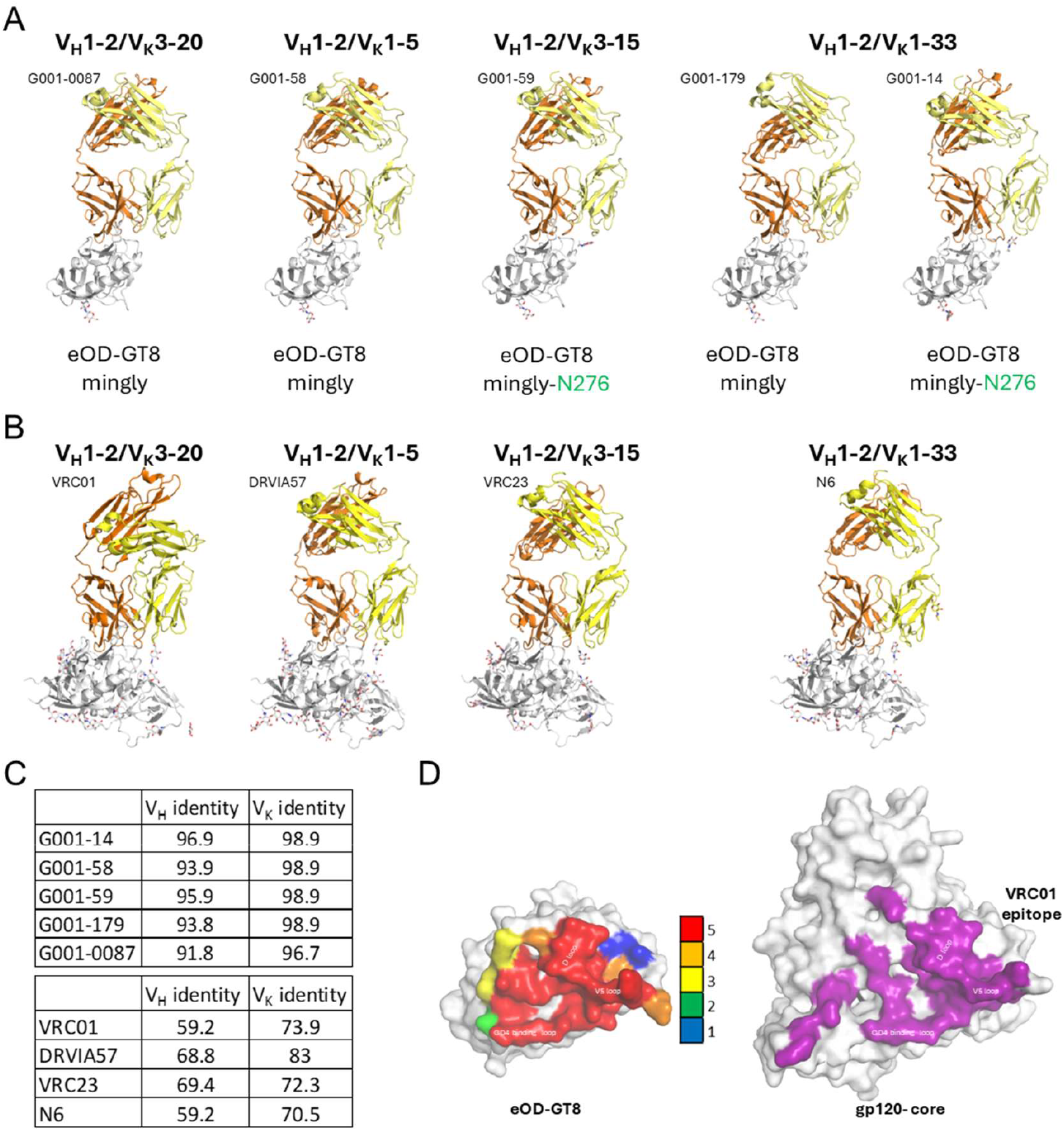
Structural characterization of G001 trial-derived antibody precursors bound to eOD-GT8. (A) X-ray crystal structures of five vaccine-elicited antibody precursors (heavy chains in orange, light chains in yellow) in complex with eOD-GT8 (gray). (B) Structures of bnAb Fab/HIV gp120 complexes encoded by four different light-chain V genes matching those found in the G001 trial-derived antibody precursors. The PDB IDs from left to right are 3NGB, 5CD5, 4J6R, and 5TE6. (C) Germline sequence amino acid identity (%) of heavy and kappa chain variable regions for precursors (top) and mature bnAbs (bottom). (D, left) Residue-specific contact frequency (heatmap) on eOD-GT8 for antibody precursors, calculated from Fab paratope interactions (color scale: 0–5 contacts/residue). (D, right) Contact residues of bnAb VRC01 (PDB: 3NGB) mapped onto HIV gp120 core, highlighting conserved CD4bs engagement. Contact residues were calculated using PDBePISA.

The five antibodies retain over 91% and 97% germline identity in the heavy and kappa chain variable regions (V_H_ and V_K_) (Fig. 1C), and converge on a shared epitope centered around the CD4bs, similar to VRC01-class bnAbs (Fig. 1D). In contrast, VRC01-class bnAbs exhibit extensive SHM, reducing their germline identity to 60% for V_H_ and 74% for V_K_ (Fig. 1C). The structures reveal that, even with minimal SHM, these antibodies effectively mimic the interaction of mature VRC01-class bnAbs with HIV Env.

To further define the shared epitope recognized by VRC01-class bnAb precursors, we utilized a paratope residue analysis. A residue-specific contact heat map illustrates a focused response to the CD4bs concentrated on CD4 binding loop, V5 loop, and D loop (Fig. 1D, Fig. S1). Notably, all five antibodies block the CD4bs by forming beta-sheet interactions with the CD4 binding loop through heavy-chain complementarity-determining region 2 (HCDR2), a key characteristic of VRC01-class bnAbs (39). This shared binding mode reinforces the critical role of HCDR2 and FR3 residues in epitope engagement and highlights the structural basis for the germline-targeting immunogen design strategy.

### Role of IGHV1-2 in VRC01-class antibody recognition of the CD4bs

IGHV1-2 plays a crucial role in CD4bs recognition (39). Previous studies have identified a defining characteristic of VRC01-class antibodies as their ability to engage the HIV-1 CD4bs predominantly through residues in the heavy-chain HCDR2 and framework region (FR) 3, with minimal reliance on HCDR3 (11, 33, 39, 47). Structural analyses of the five VRC01-class bnAb precursors reveal that they utilize the IGHV1-2-derived HCDR2 and FR3 regions for epitope recognition (Fig. 2A), mirroring the recognition in mature VRC01-class bnAbs (4). This conserved binding mechanism is further corroborated by the observation that these antibodies, as exemplified by G001-0087, display similar buried surface areas (BSAs) and hydrogen bond patterns for HCDR2 and FR3 with the CD4bs (Fig. 2B,C), underscoring the conserved nature of these interactions across the maturation process.

**Fig. 2.**
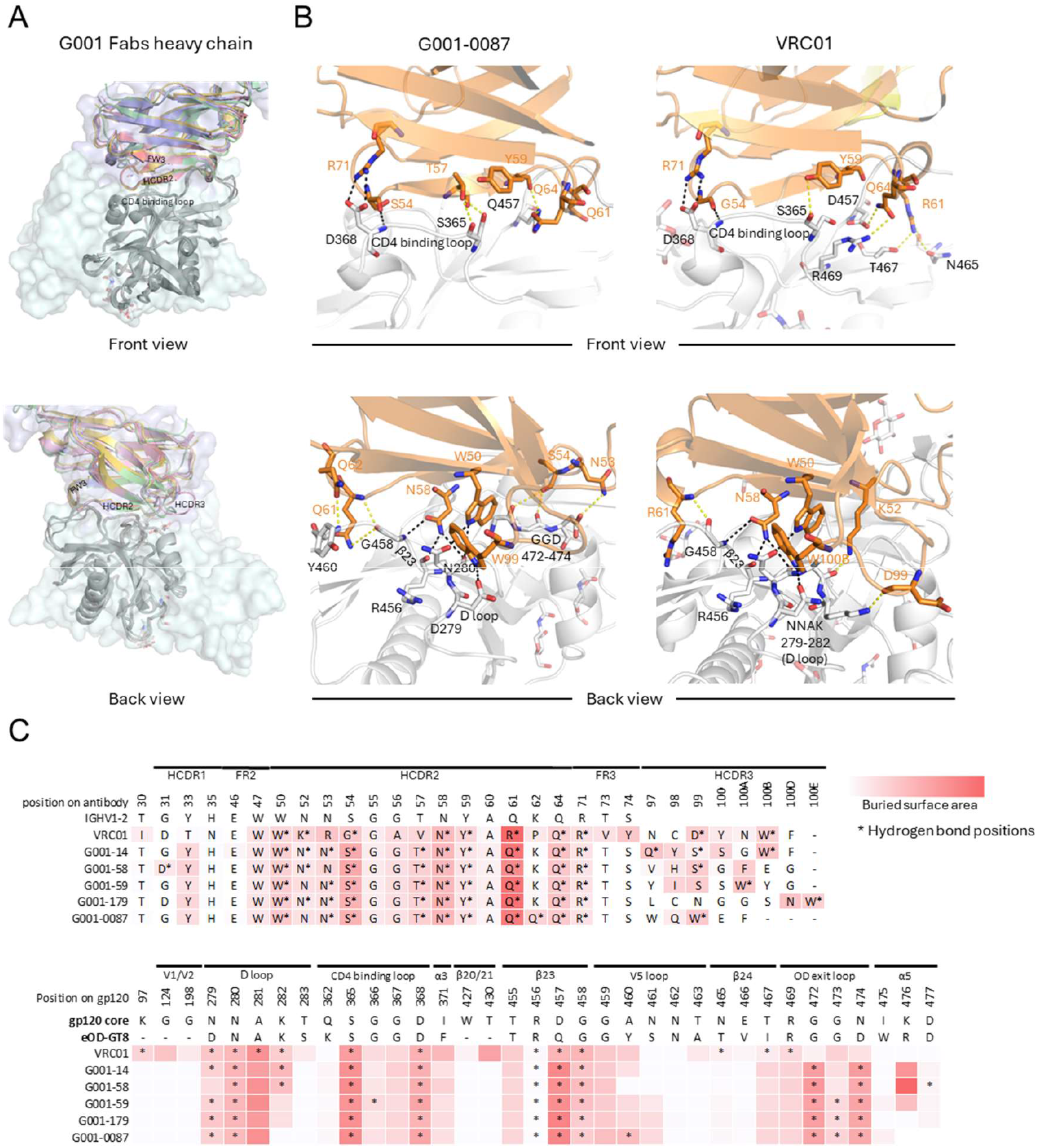
Conserved heavy chain mediated recognition of the CD4-binding site. (A) Structural alignment of bnAb VRC01 (PDB: 3NGB) and precursor heavy chains (colored: VRC01 [lightblue surface], G001-0087 [red], G001-58 [yellow], G001-59 [blue], G001-179 [pink], G001-14 [green]), performed by superimposing the eOD-GT8 antigen (gray) onto HIV gp120 core (lightcyan surface), highlighting conserved IGHV1-2 interactions. (B) Zoomed-in view of the IGHV1-2 interaction of G001-0087 with bnAb VRC01 as a comparison. Residues involved in hydrogen bonds are shown in sticks, and conserved hydrogen bonds in both antibody precursor and bnAb VRC01 are indicated by black dashed lines. All other hydrogen bonds are marked by yellow dashed lines. (C) Antibody (top) and antigen (bottom) contact residues involved in the interaction are compared in sequence, with hydrogen bonds indicated by stars and surface buried area in a red scale. Interfaces were calculated using PDBePISA.

Key interacting residues, such as W50, N58, Y59, Q64, and R71, remain unchanged from the IGHV1-2*02 germline, for bnAb VRC01 and the VRC01-class bnAb precursors studied here (Fig. 2B,C top, Fig. S2), reflecting a highly conserved mode of epitope recognition. These residues are central to forming hydrogen bonds and salt bridges that enhance binding and specificity to the HIV-1 CD4bs. In contrast, somatic mutations in bnAb VRC01, such as N52K, N53R, S54G, T57V, and Q61R, introduce slight variations in hydrogen bonding that may fine-tune binding affinity without disrupting the overall mode of recognition. While HCDR3 interactions are less conserved and exhibit more variability among these antibodies, most VRC01-class antibodies maintain a critical tryptophan residue, located five residues before the end of HCDR3 (Trp100b in VRC01, Trp99 in G001-0087, Trp100b in G001-14, Trp100a in G001-59) that forms a hydrogen bond with D/N279 of the antigen (19, 33). Interestingly, G001-58 deviates from this trend, as its phenylalanine residue (Phe100a) mediates interaction without forming a hydrogen bond.

On the epitope side, the BSAs and hydrogen bond patterns of both mature VRC01 and the bnAb precursors converge in key regions (Fig. 2C bottom). All are anchored to the CD4-binding loop (hydrogen bonds and salt bridge with S365 and D368), D loop (hydrogen bonds with D/N279 and N280), and β23 (hydrogen bonds with R456, D/Q457, and G458). Differences arise at β24 and the OD exit loop, where germline residues N53 and S54 on VRC01-class bnAb precursors preferentially form hydrogen bond with G472, G473, and D474. Conversely, bnAb VRC01 uses R61 and Q64 to form hydrogen bonds with N465, T467, and R469. These observed differences might result from structural distinctions between the eOD-GT8 and native gp120. The conserved interactions enable VRC01-class bnAb precursors to bind the epitope in a bnAb-mimicking manner while maintaining potential for further affinity maturation.

Despite differences in overall sequence identity, VRC01-class bnAbs and the vaccine-induced VRC01-class bnAb precursors share a substantial degree of similarity in their heavy-chain interactions with the CD4bs. Structural analyses highlight conserved hydrogen bonding interactions, such as the salt bridge between D368 (antigen) and R71 (antibody) and hydrogen bonds between D368 (antigen) and S/G54 (antibody). Additional critical interactions include hydrogen bonds between D/N279, N280, G458 (antigen) and N58 (antibody), between N280 (antigen) and W50 (antibody), and between D279 (antigen) and the conserved tryptophan in HCDR3 (Fig. 2B, Fig. S2). These structural studies of eOD-GT8-bound precursors show that their interactions with the immunogen mimic the binding of mature VRC01-class bnAbs to native HIV Env, reinforcing eOD-GT8 as a germline-targeting immunogen.

### Light-chain flexibility in VRC01-class bnAb precursors

Diverse but restricted light-chain V gene usage is a hallmark of both VRC01-class bnAbs and their bnAb precursors, contributing to their functional adaptability (19). In this study, monoclonal antibodies characterized by structural analyses were representative of the diverse light-chain V gene usage by VRC01-class Abs in the G001 trial. The five Abs used IGKV3-20, IGKV1-33, IGKV1-5, and IGKV3-15 (Fig. 3*A*), which were the most common light chains among VRC01-class bnAb precursors in the G001 trial and are among the most frequently used genes that encode mature VRC01-class bnAbs (48). Variations in LCDR1 conformation arise due to differences in light-chain V gene usage (Fig. 3*B*), reflecting the inherent diversity of the germline repertoire. However, structural overlay of five vaccine-induced bnAb precursors reveals a high degree of conservation in LCDR3 sequences (Fig. 3B,D,E).

**Fig. 3.**
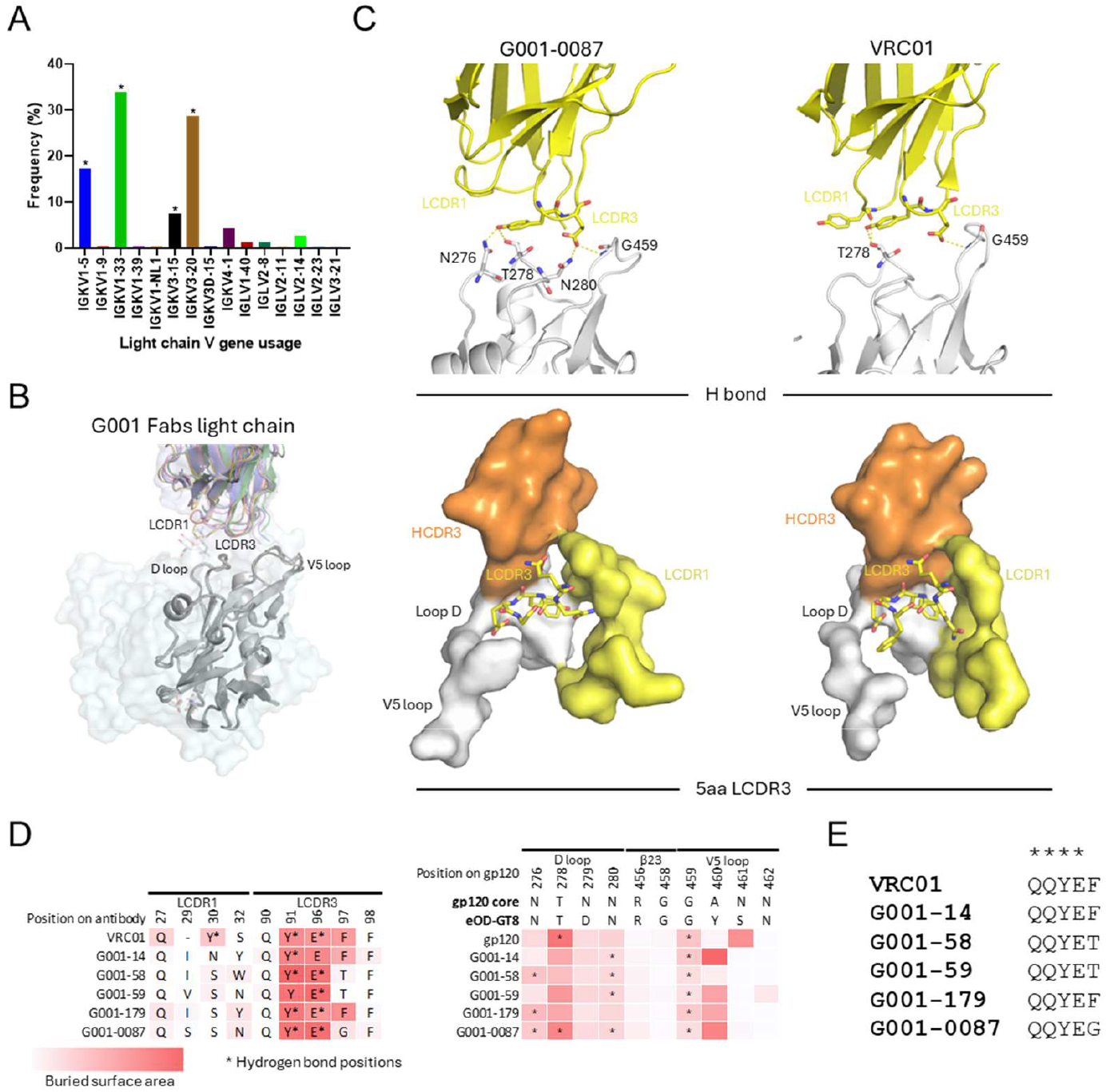
Light-chain diversity and structural constraints in antibody precursors. (A) Frequency of light-chain V gene usage among G001-derived VRC01-class bnAb precursors (n=230). Asterisks highlight the light chains used by precursors in structural analyses. (B) Structural alignment of bnAb VRC01 (PDB: 3NGB) and precursor light chains (colored as in Fig. 2A), performed by superimposing eOD-GT8 (gray) onto HIV gp120 core (lightcyan surface), showing divergent LCDR1 conformations driven by V gene usage. (C) Zoomed-in view of the light-chain interaction of G001-0087, with bnAb VRC01 as a comparison. (top: hydrogen bonds; bottom: steric constraints imposed by antigen-proximal regions). (D) Antigen (left) and antibody (right) contact residues involved in interaction are compared in sequence, with hydrogen bonds in stars and surface buried area in a red scale. Interfaces were calculated using PDBePISA. (E) LCDR3 sequence alignment of precursors and VRC01, highlighting conserved motifs (stars).

Despite differences in V gene usage, the light chain consistently contributes to the stabilization of the antigen-antibody complex by providing additional contacts that enhance binding without dominating the interaction (19, 33). BSA analyses reveal that LCDR1 and LCDR3 interact with the D loop and V5 loop of the antigen, with 2 to 3 hydrogen bonds typically forming between LCDR3 and these loops (Fig. 3C, D, Fig. S3). Notably, variations in BSA at LCDR1 are directly linked to differences in V gene usage, further highlighting the structural flexibility of the light chain. In the mature VRC01 bnAb, deletions in LCDR1 alter its conformation, enabling formation of an additional hydrogen bond with the D loop, underscoring the adaptive potential of the light chain during affinity maturation.

The five-amino-acid LCDR3 is a defining feature of VRC01-class bnAbs and their precursors (19, 33). Its short length is required by the spatial constraints within the region, where it interacts with the HCDR3, D loop, V5 loop, and LCDR1 (Fig. 3C bottom and Fig. S3). A longer LCDR3 would result in steric clashes with one or more of these loops, compromising the binding interface and reducing affinity (49). Despite variability in light-chain V gene usage, the LCDR3 sequence consistently converges to a conserved motif as previously reported (19), coinciding with the QQYEX sequence found in the crystal structures (Fig. 3E). This conservation underscores the importance of specific structural features for maintaining epitope recognition and functional compatibility across the maturation process.

### VRC01-class bnAb precursor/eOD-GT8-mingly-N276 complex structures reveal glycan adaptation

A key feature of VRC01-class bnAbs is the ability to accommodate the heavily glycosylated HIV Env surface, particularly the conserved N276 glycan. While many mature VRC01-class bnAbs achieve this by incorporating deletions or flexible residues in their LCDR1 loops (11, 33, 50), the G001 trial-derived VRC01-class bnAb precursors remain near-germline and generally lack such characteristics (19). In this study, two of the precursor complex structures (G001-59 and G001-14 in complex with eOD-GT8-mingly-N276) reveal different mechanisms for accommodating the N276 glycan without requiring LCDR1 mutations (Fig. 4A). Unlike mature bnAbs, the precursors have an intact germline LCDR1 with the glycan positioned between LCDR1 and HCDR3.

**Fig. 4.**
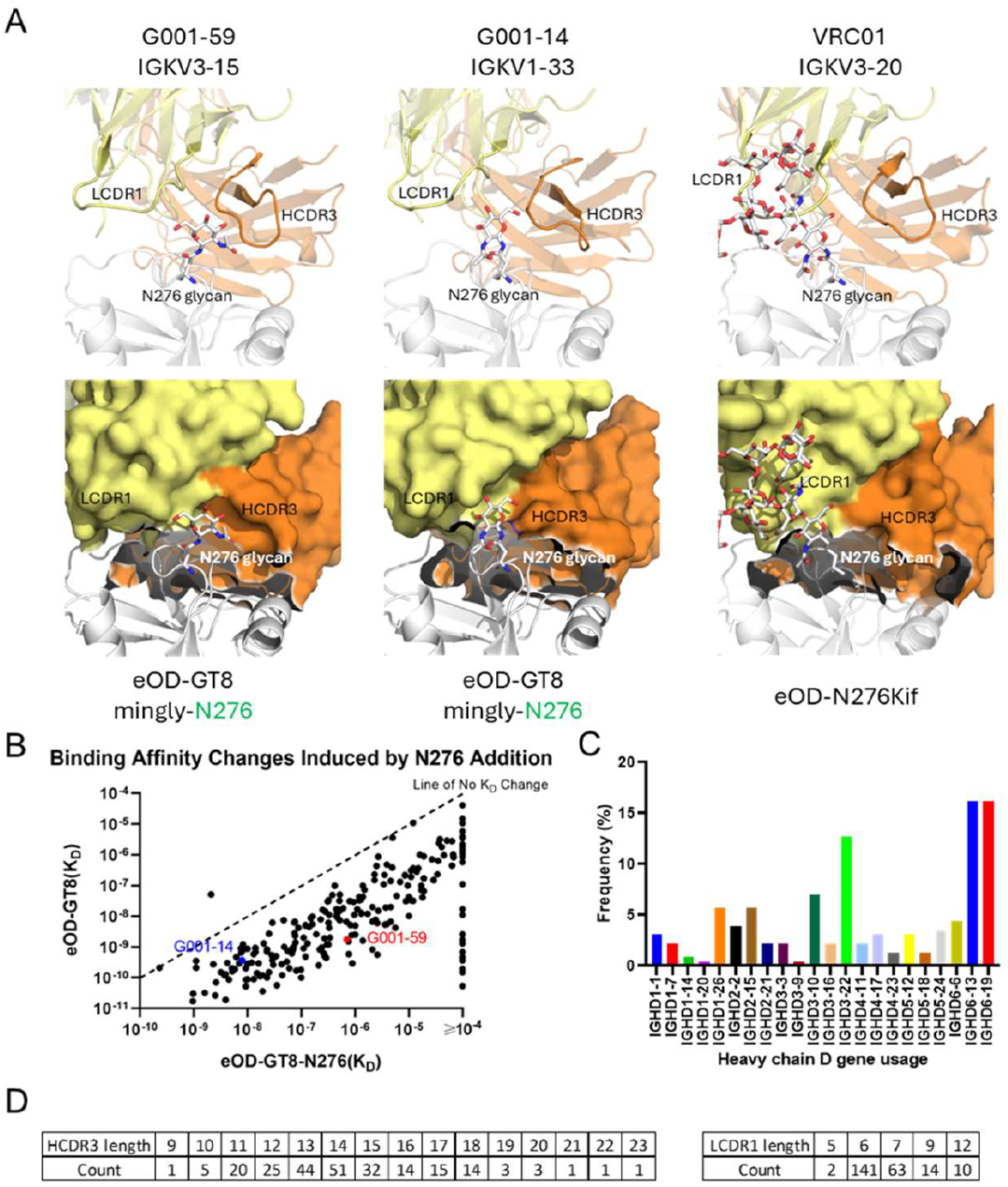
Differential accommodation of the N276 glycan in antibody precursors. (A) X-ray crystal structure comparison of how VRC01-class bnAb precursors G001-59 (left) and G001-14 (middle) engage the N276 glycan in complex with eOD-GT8-N276 (gray), compared to mature bnAb VRC01 (right) bound to a high-mannose glycan variant of eOD (PDB: 5KZC). Heavy chains are colored in orange and light chains are colored in yellow. In the cartoon view (top row), the N276 glycan is positioned between the LCDR1 and HCDR3 loops in precursors, whereas VRC01 accommodates the N276 glycan near LCDR1. The surface view (bottom row) highlights differences in specific availability for possible N276 glycan position. (B) Binding affinities (K_D_) of antibody precursors (n=230) for eOD-GT8-N276 (x-axis) and eOD-GT8 (y-axis, previously published in (19)). Most data points fall below the line of equality (dashed line), which indicates no change in K_D_. The geometric mean difference among binders to both antigens corresponds to an approximately 58-fold reduction in affinity in the presence of the N276 glycan. (C) The frequency of heavy chain D gene usage among trial-derived precursors (n=230) reflects diverse HCDR3 sequences. (D) Distribution of HCDR3 and LCDR1 lengths among trial-derived precursors (n=230), demonstrating substantial variability.

In G001-59, curvature in HCDR3 creates extra space between LCDR1 and HCDR3, allowing the N276 glycan to fit without steric clashes. In contrast, in G001-14, a slight shift in LCDR1 generates a pocket-like space between LCDR1 and HCDR3, enabling glycan accommodation. These findings demonstrate that the existence of the N276 glycan does not preclude binding to VRC01-class bnAb precursors. Instead, they highlight the potential for glycan adaptation during early stages of maturation, with further refinement likely to occur through SHM and affinity maturation.

Despite the absence of the N276 glycan in the eOD-GT8 immunogen, over 87% of antibodies tested from trial participants retained the ability to accommodate the glycan on eOD-GT8-N276 (Fig. 4B). Among 230 monoclonal antibodies analyzed, only 29 lost detectable binding upon addition of the N276 glycan to eOD-GT8. The remaining 201 demonstrated an approximate 58-fold reduction in binding affinity (K_D_) but retained a median K_D_ of 3.20×10^7^ M for eOD-GT8-N276, suggesting that glycan accommodation is achievable at the germline priming level, even in the absence of the glycan in the priming immunogen (Fig. 4B). Sequence analysis further suggests that glycan adaptation may not be strictly mediated by specific pairings of LCDR1 and HCDR3, given the substantial variation observed in heavy-chain D gene and light-chain V gene usage across the antibodies studied (Fig. 3A, Fig. 4C). This genetic variability is consistent with the observed diversity in HCDR3 and LCDR1 lengths: HCDR3 lengths ranged broadly from 9 to 23 amino acids, with the majority falling between 11 and 18 residues, whereas LCDR1 lengths were predominantly 6 or 7 amino acids, with less frequent occurrences at lengths of 5, 9, and 12 residues. The variability in those parameters indicates that glycan accommodation by early-stage precursors is not strongly constrained by specific LCDR1-HCDR3 pairings. Rather, it highlights inherent structural flexibility in diverse VRC01-class bnAb precursors, which facilitates adaptation to the N276 glycan and may subsequent maturation toward potent broadly neutralizing antibodies.

## Discussion

The development of germline-targeting vaccines represents a paradigm shift in HIV vaccine design, aiming to overcome the challenges of eliciting bnAbs through strategic activation and maturation of rare bnAb-precursor B cells (19, 35). Here, structural characterization of antibodies elicited by the eOD-GT8 60mer nanoparticle in the IAVI G001 clinical trial provides critical insights into the molecular mechanisms underlying bnAb-precursor engagement, epitope convergence, and adaptability to viral glycans. These findings provide further validation for the germline-targeting strategy and offer actionable principles for guiding booster immunogen design.

Our structural analyses reveal that vaccine-elicited VRC01-class bnAb precursors, despite retaining >90% identity to V_H_ and V_K_ germline sequences, adopt binding modes strikingly similar to those of mature VRC01-class bnAbs. The conserved engagement of the CD4bs via HCDR2 and FR3 residues demonstrates that key interactions required for bnAb function are hardwired into the IGHV1-2 germline repertoire. This structural mimicry, even in the absence of extensive SHM, underscores the precision of eOD-GT8 in selecting for precursors with bnAb-like paratopes. Notably, the antiparallel HCDR2 loop conformation, a hallmark of VRC01-class bnAbs, is recapitulated in G001 trial-derived VRC01-class bnAb precursors, reinforcing the role of germline-encoded topology in dictating epitope specificity. These observations align with prior studies showing that IGHV1-2 is critical for VRC01-class recognition of the CD4bs but extend this understanding by demonstrating that bnAb precursors elicited by human subjects can also achieve functional mimicry of mature bnAbs through conservation of key elements of the VRC01-class heavy-chain architecture (9-12, 19, 26, 51, 52).

While the heavy chain dominates CD4bs recognition by VRC01-class antibodies, light-chain diversity emerges as a critical facilitator of antigen binding (33). The use of diverse IGKV genes among trial-derived VRC01-class antibodies highlights the flexibility of the germline repertoire to employ distinct light chain frameworks while maintaining a conserved CD4bs-focused response. The structural data reveal that the light chains contribute supplementary contacts to the D and V5 loops. Importantly, the conserved five-residue LCDR3 motif, a spatial requirement to avoid steric clashes with HIV gp120, suggests that light-chain flexibility is constrained by functional necessity, ensuring compatibility with the heavy-chain-dominated paratope.

The antibodies selected for structural analysis exhibited diverse light-chain usage and were isolated from multiple individuals at different timepoints, including plasmablasts, memory B cells, and GC B cells. Although we solved only five structures, each exhibited a highly conserved, HCDR2-centered mode of binding to the HIV CD4-binding site. Accordingly, we expect that other antibodies that fulfill the VRC01-class sequence criteria and demonstrate CD4bs specificity in standard binding assays (e.g., SPR, ELISA) will similarly engage the CD4bs via this same HCDR2-dominant interaction.

Another central finding of this study is that VRC01-class bnAb precursors induced by eOD-GT8 60mer vaccination in humans have capacity to accommodate the N276 glycan, a major barrier in HIV vaccine design (53). Our structural analyses suggest that the N276 glycan can be accommodated without requiring LCDR1 deletions or hypermutations observed in mature bnAbs. While the presence of the N276 glycan reduced binding affinity by approximately 58-fold, the retention of binding in 87% of clones suggests that accommodation of the N276 glycan can be initiated at the germline level. Our findings demonstrate the feasibility of priming precursors capable of evolving toward glycan-accommodating bnAbs. However, the observed affinity reduction underscores the need for booster immunogens that selectively shepherd precursors to refine their interactions with the N276 glycan, potentially through incremental introduction of native-like glycosylation.

The modest degree of SHM of G001 trial-derived VRC01-class bnAb precursors (<10% amino acid SHM) positions them as early intermediates in the VRC01-class bnAb maturation trajectory. Their structural convergence with mature bnAbs, despite only modest mutation, suggests that critical bnAb features are established during initial priming, with subsequent maturation focusing on affinity refinement rather than epitope reconfiguration. This observation supports a “prime-boost” strategy in which initial germline-targeting immunogens lock in CD4bs specificity, while boosters progressively remove germline-targeting mutations and introduce native Env features (e.g., trimer steric environment, glycan complexity, conformational dynamics) to steer SHM toward neutralizing breadth (4, 13, 38).

While this study provides structural confirmation of successful priming of VRC01-class bnAb precursors, several questions remain. First, while the study highlights early glycan accommodation in the elicited antibodies, the engineered immunogen lacks the dense glycan shield, structural dynamics and neighboring protomers of native-like HIV trimers. Additional SHM may be required to overcome steric and electrostatic challenges posed by fully glycosylated, trimeric Env, a process not yet demonstrated in humans. Second, the study focuses exclusively on priming; the capacity of these precursors to undergo iterative affinity maturation remains to be tested in clinical settings.

The IAVI G001 clinical trial represents a proof of concept for germline-targeting in humans, and structural elucidation here of vaccine-elicited antibodies provides guidance for iterative immunogen design (19, 35). By demonstrating that VRC01-class bnAb precursors can engage the CD4bs with a bnAb-like binding mode and initiate glycan accommodation, eOD-GT8 is further validated as a foundational priming immunogen. Future boosters incorporating native-like glycans and conformational variants of HIV Env can be strategically designed to shepherd these precursors along maturation pathways observed in natural HIV infection, bridging the gap between initial priming and broad neutralization.

## Materials and Methods

### eOD-GT8 expression and purification

eOD-GT8-mingly-noN76 and eOD-GT8-mingly-N276 were transiently expressed in Expi293S cells with Expi293 Expression System Kit (Thermo Fisher Scientific), following the manufacturer’s protocol. Cell cultures were incubated at controlled condition (37°C, 8% CO2, 125 rpm) for 7 days. The supernatant was harvested, and proteins were purified using Ni-nitrilotriacetic acid (Ni-NTA) resin (Qiagen) and buffer-exchanged into Tris-buffered saline (TBS, pH 7.4). The mingly-eOD-GT8 proteins were further purified by size exclusion chromatography using HiLoad S200 16/600 column (GE Healthcare). His-tagged eOD-GT8-N276 used in SPR was produced by transient transfection of HEK-293F cells and purified as previously described (30).

### Fab expression and purification

Fabs were transiently expressed in Expi293F cells using a 3:2 plasmid ratio (heavy chain:light chain) with the Expi293 Expression System Kit (Thermo Fisher Scientific), following the manufacturer’s protocol. Cell cultures were incubated at controlled condition (37°C, 8% CO2, 125 rpm) for 7 days. The supernatant was harvested, and Fab proteins were purified using CaptureSelect CH1-XL Affinity Matrix (Thermo Fisher Scientific). Eluted Fabs were dialyzed into TBS and further purified by size-exclusion chromatography on an HiLoad S200 16/600 column (GE Healthcare).

### Sample preparation and crystallization

Purified Fab/eOD-GT8 complexes were prepared by mixing Fab and eOD-GT8 at a 1:1.2 molar ratio and incubating at 4°C overnight. Complexes were further purified by size-exclusion chromatography on a HiLoad S200 16/600 column in TBS buffer. The purified complexes were concentrated to 12 mg/mL and used for crystallization screening. Screening plates were set up on our robotic CrystalMation system (Rigaku) at The Scripps Research Institute using the vapor diffusion method in sitting drops containing 0.1 μL of protein and 0.1 μL of reservoir solution.

### X-ray structure determination and refinement

Crystals of eOD-GT8 variants and G001-series Fabs were produced under the following conditions. Crystals of eOD-GT8-mingly + G001-0087-Fab were obtained in 20% (w/v) PEG 3000, 0.1 M sodium citrate pH 5.5, with addition of 10% (v/v) ethylene glycol, which was increased to 25% (v/v) as cryoprotectant before flash-cooling in liquid nitrogen. The eOD-GT8-mingly + G001-58-Fab crystals grew in 20% (w/v) PEG 3350 and 0.2 M magnesium chloride at pH 5.8, with 15% (v/v) ethylene glycol added for cryoprotection. The eOD-GT8-mingly-N276 + G001-59-Fab crystals grew in 20% (w/v) PEG 3350 and 0.2 M magnesium sulfate at pH 5.9, with 10% (v/v) ethylene glycol added as cryoprotectant. The eOD-GT8-mingly + G001-179-Fab crystals grew in 20% (w/v) PEG 3350 and 0.2 M ammonium sulfate at pH 6.0, with 15% (v/v) ethylene glycol added as cryoprotectant. Finally, the eOD-GT8-mingly-N276 + G001-14-Fab crystals grew in 10% (w/v) PEG 8000, 0.2 M zinc acetate, and 0.1 M MES pH 6.0; no cryoprotectant was required.

Diffraction data for each crystal were either collected at beamline 12-1 of the Stanford Synchrotron Radiation Lightsource (SSRL) or beamline 23-ID-B at the Advanced Photon Source (APS), Argonne National Laboratory (see Table S2). All datasets were indexed, integrated, and scaled with the HKL-2000 software package. The structures were solved by molecular replacement using the program PHASER. A previously determined eOD-GT8 structure (PDB: 5IDL) was used as the search model for the eOD-GT8 component, while Fab models generated by Repertoire Builder were used as the search model for the antibody components. Iterative model building and refinement were carried out using Coot and Phenix, respectively.

### Surface Plasmon Resonance

Monoclonal IgGs from the G001 clinical trial were produced previously and assessed for affinity to eOD-GT8-N276 (not eOD-GT8-mingly-N276) using SPR as previously described (19). Antibody sequences, characteristics and SPR measurements are tabulated in Data S1.

### Data, Materials, and Software Availability

Structure factors and coordinates for antibody Fab fragments (G001-0087, G001-58, G001-59, G001-179, and G001-14) in complex with eOD-GT8-mingly or eOD-GT8-mingly-N276 have been deposited in Protein Data Bank with PDB ID 9OAO, 9OAP, 9OAQ, 9OAR and 9OAS.

## Acknowledgments

We thank Henry Tien for handling the automated robotic crystal screening at The Scripps Research Institute. This research was supported by the NIH National Institute of Allergy and Infectious Diseases (NIAID) UM1 AI144462 (Scripps Consortium for HIV/AIDS Vaccine Development) (to W.R.S., I.A.W.), by the Ragon Institute of MGH, MIT, and Harvard (to W.R.S.); by the International AIDS Vaccine Initiative (IAVI) Neutralizing Antibody Consortium (NAC) and Center (W.R.S., I.A.W.); and through the Collaboration for AIDS Vaccine Discovery for the IAVI NAC Center (W.R.S., I.A.W.). X-ray diffraction datasets were collected at the Stanford Synchrotron Radiation Lightsource (SSRL) beamline 12-1 and the Advanced Photon Source (APS) beamline 23-ID-B (GM/CA). This research used resources of the SSRL, SLAC National Accelerator Laboratory, which is supported by the U.S. Department of Energy, Office of Science, Office of Basic Energy Sciences under Contract No. DE-AC02–76SF00515. The SSRL Structural Molecular Biology Program is supported by the DOE Office of Biological and Environmental Research, and by the National Institutes of Health, National Institute of General Medical Sciences (including P41GM103393), and the Advanced Photon Source; a U.S. Department of Energy (DOE) Office of Science User Facility operated for the DOE Office of Science by Argonne National Laboratory under Contract No. DE-AC02-06CH11357.

## Author Contributions

X.L., C.A.C., W.R.S. and I.A.W. designed research; X.L., R.T., M.K. and D.L. expressed and purified proteins; X.L. crystallized proteins; X.L. and M.Y. collected x-ray data, X.L. determined, refined and analyzed crystal structures; O.K. performed surface plasmon resonance; X.L., C.A.C., W.R.S. and I.A.W. analyzed data; X.L., C.A.C., W.R.S. and I.A.W. wrote the paper; and all authors reviewed the paper.

