## Supplementary Material for "Structural insights into VRC01-class bnAb precursors with diverse light chains elicited in the IAVI G001 human vaccine trial"

**Fig. S1. Conserved CD4bs engagement by bnAbs.** Epitope defined by contact residues (as in Fig. 1D) for bnAbs DRVIA57 (PDB: 5CD5), VRC23 (PDB: 4J6R) and N6 (PDB: 5TE6) complexed with HIV gp120-core. See also Fig. 1D for VRC01.

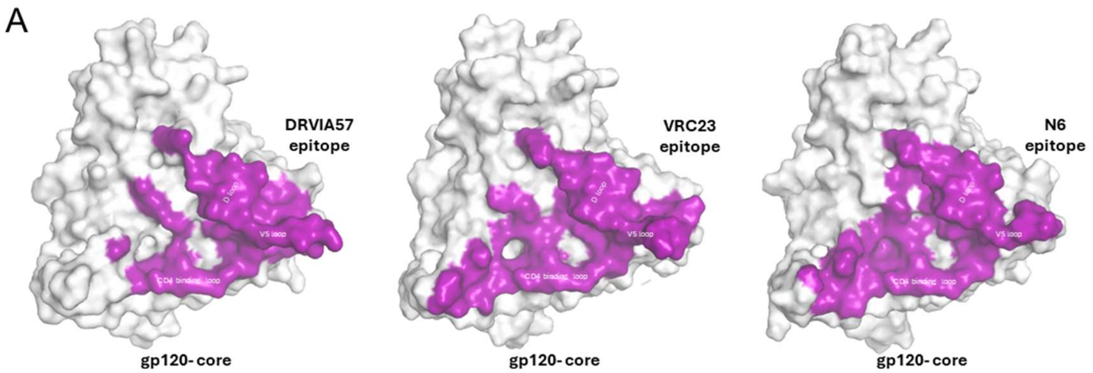

**Fig. S2. Conserved heavy-chain interactions across antibody precursors.** Zoomed-in view of the IGHV1-2 interaction (as in Fig. 2B) for four G001 antibody precursors (G001-14, G001-58, G001-59 and G001-179). Residues involved in hydrogen bonds are shown in sticks, and conserved hydrogen bonds in both antibody precursor and bnAb VRC01 are indicated by black dashed lines. All other hydrogen bonds are marked by yellow dashed lines.

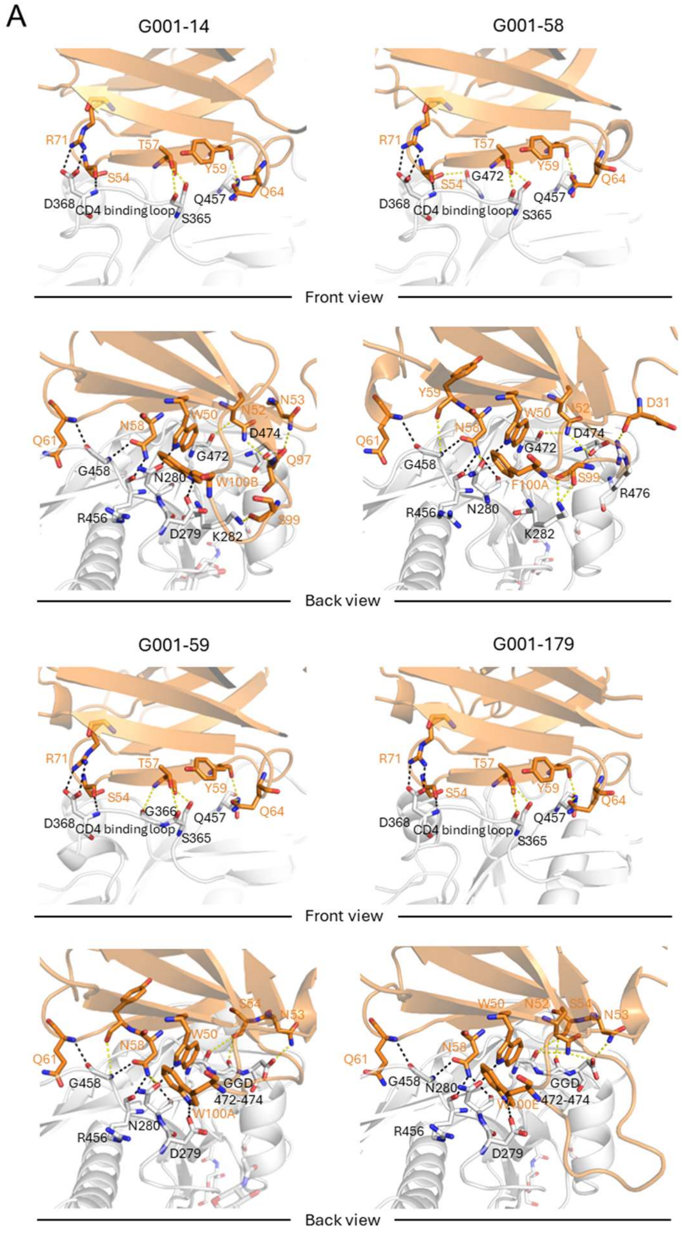

**Fig. S3. Light-chain interactions across antibody precursors.** Zoomed-in view of light-chain interactions (as in Fig. 3C) for four G001 antibody precursors. (G001-14, G001-58, G001-59 and G001-179). (top: hydrogen bonds; bottom: steric constraints imposed by antigen-proximal regions).

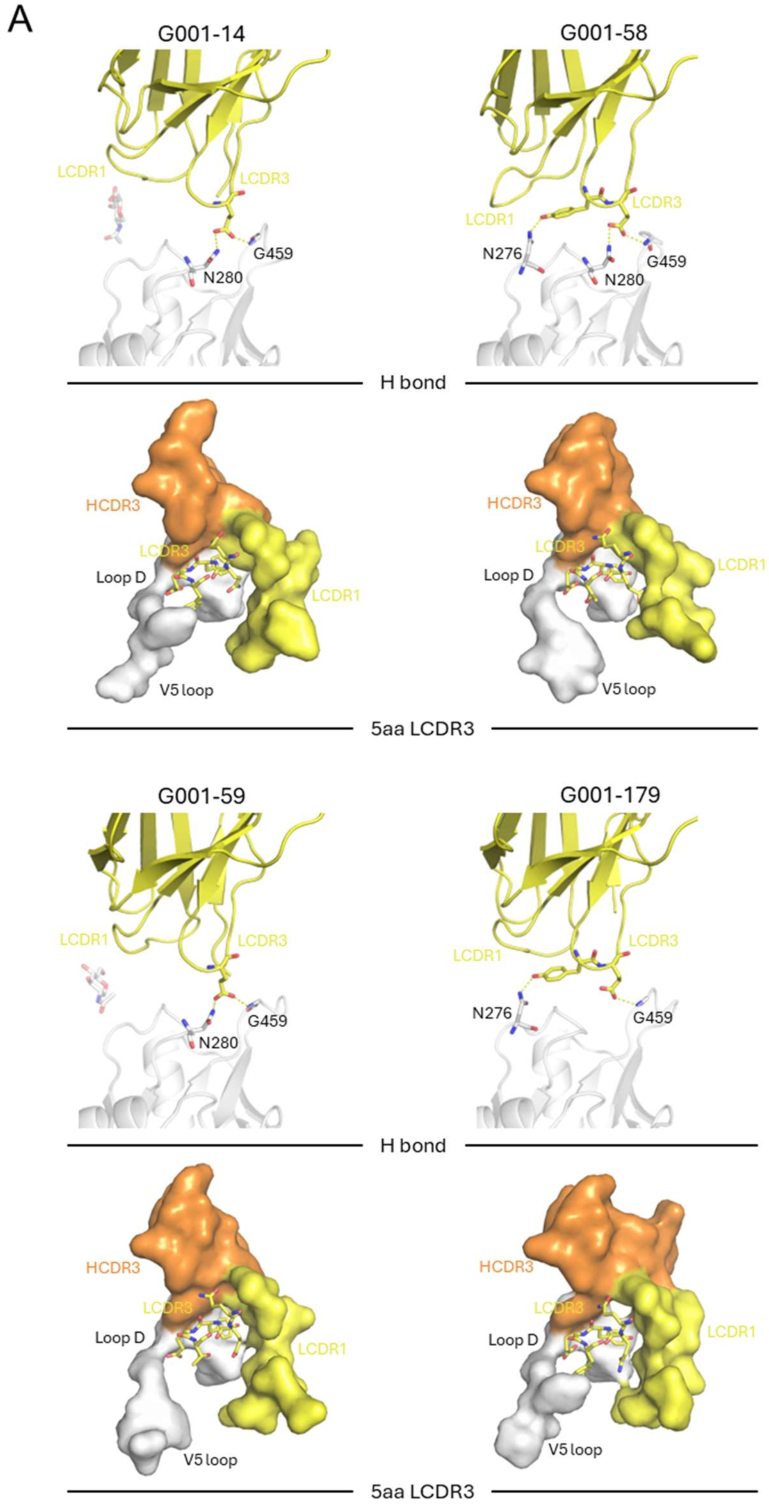

### Table S1. Properties of the five VRC01-class bnAb precursors selected for structural analysis.

| Antibody name | Timepoint | eOD-GT8 Kon | eOD-GT8 Koff | eOD-GT8 KD (M) | Specimen Type | Heavy V gene | Heavy D gene | Heavy J gene | HCDR3 length | Heavy V gene identity (aa) | Light V gene | Light J gene | HCDR3 length | Light V gene identity (aa) |
|---|---|---|---|---|---|---|---|---|---|---|---|---|---|---|
| G001_gp1_V10_VRC01c_2021-0087 | week 16 | 4.20E+05 | 6.20E-05 | 1.50E-10 | PBMC | IGHV1-2*04 | IGHD3-10*01 | IGHJ4*02 | 11 | 91.84% | IGKV3-20*01 | IGKJ4*01 | 5 | 96.70% |
| G001_LD_07A_VRC01c_58 | week 9 | 3.98E+05 | 1.74E-04 | 4.37E-10 | PB | IGHV1-2*04 | IGHD5-12*01 | IGHJ4*02 | 13 | 93.88% | IGKV1-5*03 | IGKJ1*01 | 5 | 98.90% |
| G001_LD_07A_VRC01c_59 | week 9 | 3.94E+05 | 6.93E-04 | 1.76E-09 | PB | IGHV1-2*04 | IGHD6-13*01 | IGHJ4*02 | 13 | 95.92% | IGKV3-15*01 | IGKJ1*01 | 5 | 98.90% |
| G001_gp1_V10_VRC01c_179 | week 16 | 4.50E+05 | 5.90E-05 | 1.30E-10 | PBMC | IGHV1-2*02 | IGHD2-15*01 | IGHJ4*02 | 17 | 93.81% | IGKV1-33*01 | IGKJ3*01 | 5 | 98.90% |
| G001_LD_V05_VRC01c_14 | week 3 | 2.10E+05 | 7.60E-05 | 3.70E-10 | FNA | IGHV1-2*04 | IGHD6-19*01 | IGHJ4*02 | 14 | 96.94% | IGKV1-33*01 | IGKJ4*01 | 5 | 98.90% |

### Table S2. Data collection and refinement statistics.

| **Data collection** | eOD-GT8-mingly + G001-0087-Fab | eOD-GT8-mingly + G001-58-Fab | eOD-GT8-mingly-N276 + G001-59-Fab | eOD-GT8-mingly + G001-179-Fab | eOD-GT8-mingly-N276 + G001-14-Fab |
|---|---|---|---|---|---|
| Beamline | APS 23-ID-B | SSRL 12-1 | SSRL 12-1 | SSRL 12-1 | SSRL 12-1 |
| Wavelength (Å) | 1.0332 | 0.9795 | 0.9795 | 0.9795 | 0.9795 |
| Space group | I 2 2 2 | C 2 2 $2_1$ | I 2 2 2 | P 1 | C 1 2 1 |
| Unit cell parameters | | | | | |
| a, b, c (Å) | 102.5, 136.171, 152.944 | 110.1, 174.9, 93.6 | 102.7, 134.0, 154.0 | 49.2, 65.9, 122.1 | 261.1, 75.5, 158.1 |
| α, β, γ (°) | 90, 90, 90 | 90.0, 90.0, 90.0 | 90.0, 90.0, 90.0 | 89.9, 97.0, 90.0 | 90.0, 92.4, 90.0 |
| Resolution (Å) [a] | 50.00 - 2.60 (2.64 - 2.58) | 50.00-2.80 (2.87-2.80) | 50.00-2.54 (2.58-2.54) | 50.00-1.97 (2.00-1.94) | 50.00-2.81 (2.86-2.79) |
| Unique reflections [a] | 33,387 (1,503) | 22,255 (1,109) | 34,890 (1,696) | 96,860 (4,358) | 75,138 (3,714) |
| Redundancy [a] | 10.5 (4.0) | 12.8 (12.2) | 11.4 (6.1) | 3.3 (2.8) | 6.8 (6.9) |
| Completeness (%) [a] | 99.3 (90.9) | 99.1 (99.4) | 99.0 (98.3) | 89.6 (80.8) | 98.4 (99.2) |
| $\langle I/\sigma_I \rangle$ [a] | 12.3 (1.1) | 8.3 (1.0) | 14.5 (1.0) | 5.8 (2.1) | 7.7 (1.3) |
| $R_{sym}$ [b] (%) [a] | 25.9 (115.9) | 53.7 (686.4) | 16.5 (191.4) | 26.6 (139.3) | 31.4 (346.6) |
| $R_{pim}$ [b] (%) [a] | 7.6 (49.9) | 14.9 (194.8) | 4.8 (72.5) | 14.3 (78.5) | 11.9 (131.3) |
| $CC_{1/2}$ [c] (%) [a] | 98.0 (42.9) | 98.7 (41.1) | 99.7 (39.3) | 95.0 (32.3) | 97.8 (31.1) |
| **Refinement statistics** | | | | | |
| Resolution (Å) | 47.74 - 2.58 | 47.44-2.80 | 38.50-2.54 | 26.28-1.94 | 38.18-2.79 |
| Reflections (work) | 33,110 | 21,156 | 34,672 | 96,730 | 74,974 |
| Reflections (test) | 1,631 | 1,070 | 1,668 | 5,082 | 3,808 |
| $R_{cryst}$ [d] / $R_{free}$ [e] (%) | 23.8 / 25.3 | 21.3/24.9 | 20.8/23.0 | 21.3/24.8 | 21.4/25.4 |
| No. of atoms | 4,696 | 4,541 | 4,719 | 10,722 | 18,474 |
| Antigen | 1,288 | 1,288 | 1,284 | 2,576 | 5,136 |
| Fab | 3,228 | 3,225 | 3,259 | 6,448 | 13,036 |
| Glycan | 28 | 28 | 42 | 56 | 168 |
| Solvent | 152 | - | 134 | 1,642 | 134 |
| Average *B*-values (Å$^2$) | 48 | 63 | 58 | 23 | 67 |
| Antigen | 56 | 69 | 66 | 22 | 61 |
| Fab | 45 | 60 | 54 | 23 | 69 |
| Glycan | 73 | 96 | 94 | 31 | 88 |
| Solvent | 43 | - | 50 | 25 | 51 |
| Wilson *B*-value (Å$^2$) | 44 | 64 | 56 | 18 | 61 |
| **RMSD from ideal geometry** | | | | | |
| Bond length (Å) | 0.003 | 0.003 | 0.002 | 0.007 | 0.004 |
| Bond angle (°) | 0.55 | 0.50 | 0.54 | 0.92 | 0.71 |
| **Ramachandran statistics (%)** | | | | | |
| Favored | 96.4 | 95.7 | 97.8 | 96.6 | 96.6 |
| Outliers | 0.00 | 0.34 | 0.17 | 0.17 | 0.13 |
| **PDB ID** | 9OAO | 9OAP | 9OAQ | 9OAR | 9OAS |
